# *In silico* bacteria evolve robust cooperation via complex quorum-sensing strategies

**DOI:** 10.1101/598508

**Authors:** Yifei Wang, Jennifer B. Rattray, Stephen A. Thomas, James Gurney, Sam P. Brown

## Abstract

Many species of bacteria collectively sense and respond to their social and physical environment via ‘quorum sensing’ (QS), a communication system controlling extracellular cooperative traits. Despite detailed understanding of the mechanisms of signal production and response, there remains considerable debate over the functional role(s) of QS: in short, what is it for? Experimental studies have found support for diverse functional roles: density sensing, mass-transfer sensing, genotype sensing, etc. While consistent with theory, these results cannot separate whether these functions were drivers of QS adaption, or simply artifacts or ‘spandrels’ of systems shaped by distinct ecological pressures. The challenge of separating spandrels from drivers of adaptation is particularly hard to address using extant bacterial species with poorly understood current ecologies (let alone their ecological histories). To understand the relationship between environmental challenges and trajectories of QS evolution, we used an agent-based simulation modeling approach. Given genetic mixing, our simulations produce behaviors that recapitulate features of diverse microbial QS systems, including coercive (high signal / low response) and generalized reciprocity (signal auto-regulation) strategists — that separately and in combination contribute to QS-dependent resilience of QS-controlled cooperation in the face of diverse cheats. We contrast our *in silico* results with bacterial QS architectures that have evolved under largely unknown ecological contexts, highlighting the critical role of genetic constraints in shaping the shorter term (experimental evolution) dynamics of QS. More broadly, we see experimental evolution of digital organisms as a complementary tool in the search to understand the emergence of complex QS architectures and functions.

**Author summary:** Bacteria communicate and cooperate using complex cell-cell signaling systems known as quorum-sensing (QS). While the molecular mechanisms are often well understood, the reasons why bacteria use QS are less clear — how has QS aided survival and growth? The answer to this question is dependent on the environment of adaptation, and unfortunately our current understanding of QS bacterial ecology is broadly lacking. To address this gap, we studied the evolution of ‘digital organisms’, individual-based computer simulations of bacterial populations evolving under defined environmental contexts. Our results pinpoint how simple environmental challenges (variable density and genetic mixing) can lead to the emergence of complex strategies that recapitulate features of bacterial QS, and open a path towards reverse-engineering the environmental drivers of QS.

## Introduction

Many species of bacteria are highly social, investing in the secretion of multiple costly molecules in order to gain collective benefits. The benefits of microbial collective action are diverse, including extracellular digestion of complex molecules (via secretion of exo-enzymes [1]), access to limiting iron (via secretion of siderophores [2–4]), construction of defensive biofilms (via secretion of exopolysaccharide building blocks [5, 6] or anti-competitor toxins (via secretion of antibiotics and bacteriocins [7–10]).

Despite the shopping list of potential collective benefits, microbes like other social organisms face the challenge of identifying under what conditions costly investment in collective activity is going to return a net benefit. A striking commonality across multiple species of bacteria is the use of complex cell-cell signaling mechanisms known as quorum-sensing (QS) to control and coordinate the expression of social behaviors mediated by secreted factors [11]. Quorum-sensing bacteria secrete diffusible signal molecules, and respond to the accumulation of signal in their environment by changing global gene expression — increasing the relative rate of production of costly exo-enzymes, toxins and other secreted factors [12–16].

The QS control of secreted factors has been long argued to allow bacteria to limit their costly investments in collective action to environments where a focal strain of bacteria are at high densities (‘quorate’) and therefore able to effectively modify their environment [17, 18], and it has more recently been demonstrated that cooperative exo-enzyme production is indeed more beneficial at higher densities [19]. Other theories suggest that QS allows bacteria to sense their physical environment such as the degree of containment or mass transfer [20, 21], the biochemical environment [22, 23] or through the use of multiple signals to simultaneously resolve both physical environment and population density [24].

Bacteria also face potential uncertainty over the genotypic mix of their environment — are individuals exploiting a local environment as a single clone, or in the context of other strains or species? It is established that QS-controlled cooperative behaviors such as exo-enzyme production are vulnerable to social exploitation by ‘cheat’ strains that can reap the rewards without paying the costs of secretions, and that this vulnerability can be mitigated by positive genetic assortment / kin selection [25–28], and also by the coupling of directly beneficial traits under QS control [23, 29]. Other studies have explored how the architecture of QS allows individuals to sense and strategically respond to variations in the genotypic environment [30–33], and on a longer evolutionary time-scale how the tuning and architecture of QS is itself potentially shaped by enduring patterns of genetic conflict [16, 25, 30, 34, 35].

The complexity of QS also raises the possibility of additional dimensions of social conflict beyond the extent of costly exo-product production. In common with any communication system, QS is the product of the coupled evolution of a signal and a response strategy [36]; implying that the return on investment in signaling depends on the response strategy (signal affinity), and vice versa. This coupling highlights that to hit a defined density (or diffusion, etc) threshold there are multiple signal / response equilibria, or ‘social norms’: for example a high signaling / low response ‘shouting and deaf’ equilibrium can be equally effective for a clonal population detecting a threshold as low signaling / high response ‘conspiratorial whisper’ equilibrium [25]. These alternate equilibria are not equivalent however in a non-clonal context; a high signal / low response strategist will act to coercively induce greater cooperative returns whenever mixed with a low signal / high response strategist, leading to their predicted dominance under conditions of intermediate genetic mixing [25].

The complexity and potential multi-functionality of QS has lead to a growing number of experimental studies designed to experimentally determine the functional roles of QS across multiple bacterial systems [17, 19, 20, 24, 30, 32, 37–43]. Across these experimental systems, there is evidence that QS systems can limit cooperative investments to high density [19], low mass transfer [21, 44] and clonal [30, 32] environments — in some cases these responses have demonstrated fitness advantages [19, 30, 32]. While consistent with theoretical predictions, these results cannot separate whether these functional roles were the drivers of QS evolution, or simply fortuitous byproducts or ‘spandrels’ [45] of a complex system driven by other ecological pressures.

The challenge of separating drivers of adaptation from ‘spandrel’ properties is particularly hard to address using extant bacterial species with complex QS architectures that have evolved under ecological conditions that we scarcely understand. To address this challenge we turn to *in silico* experimental evolution [46], which allows us to evolve signaling strategies among digital organisms under defined ecological challenges. We study the joint evolution of multiple QS component traits (basal signal production, cooperative response, signal auto-regulation) under a range of conditions of environmental heterogeneity (variable densities and variable genetic mixing among groups).

### Methods Summary

In our agent-based *in silico* evolution experiments, we first consider two evolving traits, basal production rate (*p*) and signal response threshold (*S_Th_*). In subsequent simulations, we introduce an additional evolving trait, the auto-regulation ratio (*r*), the ratio of fully induced to basal signal level. These co-evolving traits combine to govern individual cell decisions to turn on or off cooperation in different environmental conditions. Populations are subject to selection based on individual cooperation payoffs (incorporating costs of cooperation and signaling, and environment-dependent benefits), and evolve for a fixed number of generations as shown in Fig. 1 (see *Appendix* for more model implementation details).

**Fig 1.**
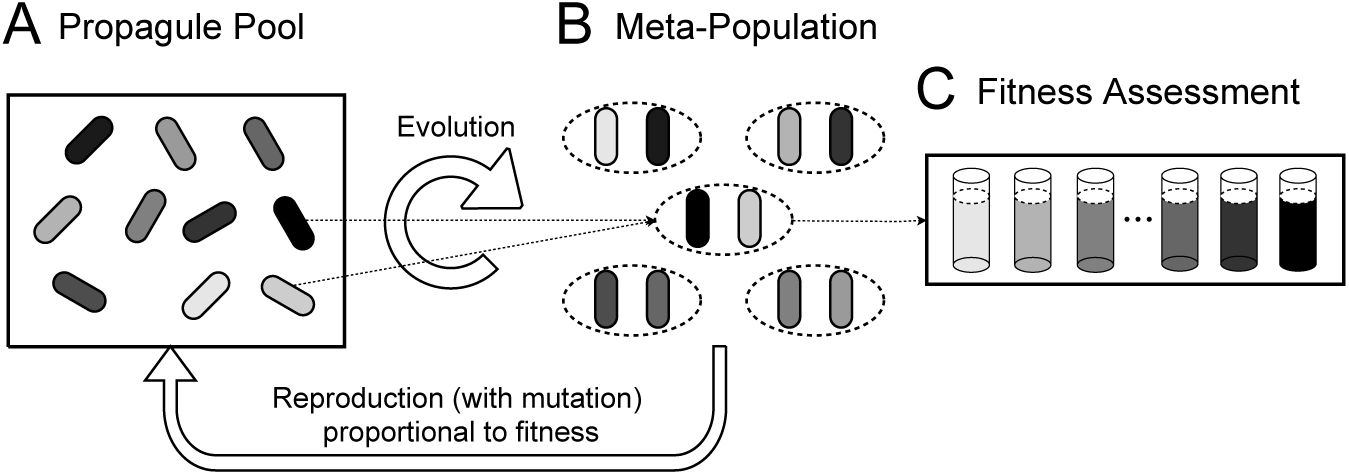
*In silico* evolution of quorum sensing. **(A)** The propagate pool at time zero is a strategically uniform population of *in silico* individuals. (B) In each generation, individuals are distributed into locally interacting sub-populations depending on the condition of genetic mixing (defined by number of founding cells per sub-population — two founders per sub-population in this illustration). (C) The fitness of all individual founders is evaluated across a range of testing environments (different population densities). Then, individuals are selected proportionally to their payoffs for clonal reproduction but are subject to mutation in their quorum sensing traits (signal production, threshold to response, and in some simulations, auto-regulation). Finally, the offspring pool with the same size as the initial propagate pool was formed for the next generation.

The fitness of each genotype in each sub-population is assessed across a spectrum of potential bacterial carrying capacities (see Fig. 1). For carrying capacity (at density *N*), signaling and resulting cooperative responses are described by an ODE governing extracellular signal concentration. Specifically, we consider the following two scenarios of signal dynamics for QS-controlled cooperation in absence (Eq. (1)) and presence (Eq. (2)) of auto-regulation, respectively.

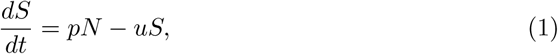

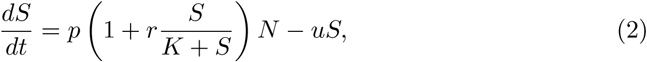

where *S* is the local signal concentration, *t* is time, *N* is the stationary phase cell density, *p* is the basal signal production rate, *r* is the ratio of auto-regulation production to basal signal production, *K* is the half concentration value, and *u* is the signal decay rate. We assume that signal concentration rapidly equilibrates to *S*^*^ (see *Appendix*), and use this value to determine individual cooperation. Individuals will turn on their cooperative phenotype only when the local signal concentration is higher than the individual response threshold (*S*^*^ > *S*_*Th*_). Both signaling and cooperation are costly to individuals, but they benefit from cooperation only when the local cellular density is above a certain threshold *N_Th_*. Therefore cost-effective cooperative behavior is dependent on the effective tuning of QS to identify an underlying density threshold. For constitutive cooperation (No QS), individuals will always turn on cooperation regardless of local signaling environment — they do not have the ability to make social informed choices. The Julia source code can be downloaded here: https://bit.ly/2u3OcSM

## Results

### The quorum sensing coordination game in a clonal context

We begin by defining a cooperative trait with a threshold density dependent benefit (see *Methods Summary* and *Appendix*), and illustrate that the joint evolution of signal production (*p*) and signal threshold to response (*S_Th_*) can tune individual cooperative behavior to solve the ‘density sensing’ problem (Fig. 2A).

**Fig 2.**
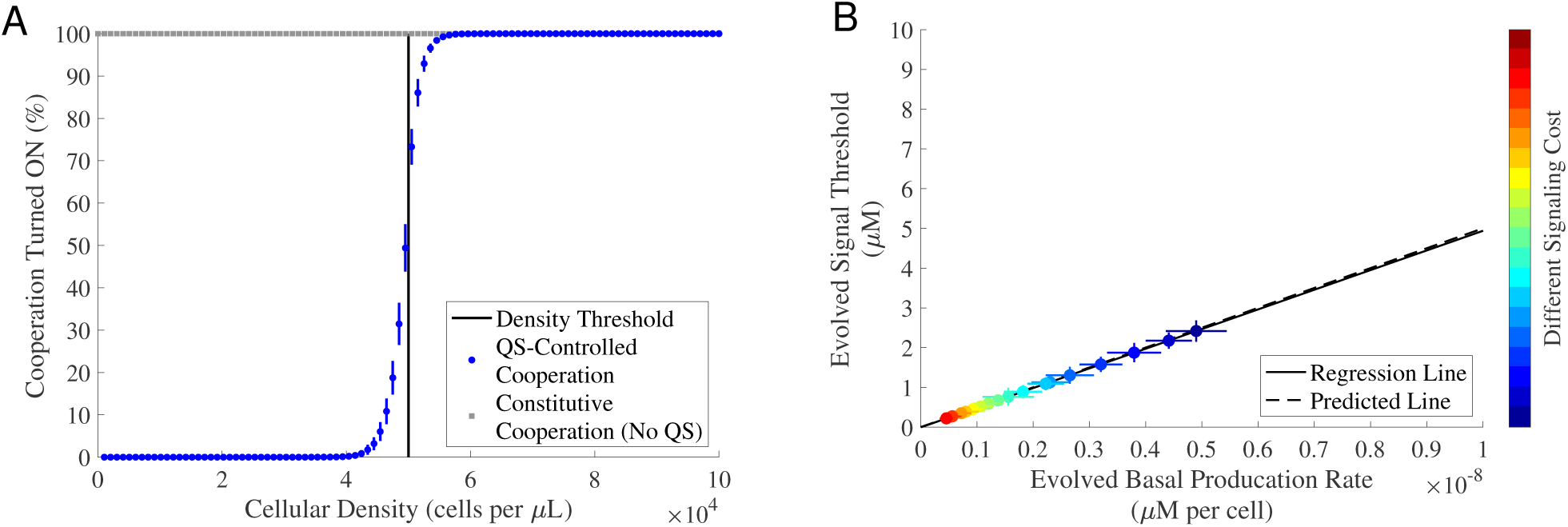
Signal production rate and threshold response co-evolve in a quorum sensing co-ordination game. We evolved 5, 000 initially identical genotypes for 5, 000 generations in a patchy, variable density environment (see Fig. 1). In all simulations, the cost of cooperation was fixed and there was no auto-regulation. (A) We used a fixed cost of signaling, and recorded the percentage of evolved individuals who turned on cooperation in 100 testing environments where the cellular density varied from 10^1.5^ to 10^5^ cells per *μL*. Note constitutive individuals, by definition, will always turn on cooperation in all environments as shown by the gray dotted line. The vertical black line indicates the pre-set density threshold N_Th_ in our simulations. (B) Varying the cost of signaling (*C*_*sig*_[0, 10^10^]; step size: 510^8^) predictably impacts the balance of signal production and response. Each dot represents the evolved mean results (averaged over the last 50 generations). The color-bars indicate different values of *C*_*sig*_ from low signaling cost (dark blue) to high singling cost (dark red). The solid black line is the regression line fitted using the generalized linear model with a normal distribution: *R*^2^ = 1, *F*-test, *p* = 5.511 × 10^−50^. The dashed black line is the predicted line calculated by *S*_*Th*_ = _*p*_*N*_*Th*_/*u*. The horizontal and vertical error bars represent the standard deviation over 30 replications. The remaining parameters used in the simulations can be found in *Appendix*, Table S1.

Our analyses show that bacteria solve this problem by resolving a co-ordination game between signal and response strategies. Given our model of extracellular signal dynamics in the absence of auto-regulation (Eq. (1), see *Methods Summary*), we can see that for a given stationary density *N*, signal concentration *S* will equilibrate to *S*^*^ = *pN* /*u*, where *u* is the rate of environmental signal degradation. Given a critical density threshold (for cooperation to pay) of *N_Th_*, we can define the critical signal threshold *S_Th_* = *pN_Th_*/*u*, so that signal will trigger cooperation if *S*^*^ > *S*_*Th*_. Together, this implies that QS bacteria will cooperate when *pN* /*u > S_Th_*. Given that both *p* and *S_Th_* are evolutionary variables, the joint evolution of both traits forms a coordination game — the optimal value of signal production *p* depends on *S_Th_*, and vice versa. In Fig. 2B, we illustrate the nature of this coordination game by varying the cost of signaling. As shown in Fig. 2B, as the cost of signaling increases, the basal signaling rate *p* evolves to a lower intensity. As a consequence, the signal threshold *S_Th_* decreases accordingly in order to maintain resolution of density threshold *N_Th_*, as predicted by *S_Th_* = *pN_Th_/u* (dashed line in Fig. 2B). In *Appendix*, Figs. S1A and S1B (also see *Appendix*, Fig. S2) we illustrate that the evolved coordination of signaling *p* and response *S_Th_* is also dependent on environmental factors governing signal decay rate and noise. Together, these results illustrate that clonal bacteria can coordinate their behaviors to adjust their signal production rate and their responsiveness to signal, and this QS-controlled cooperation is superior to constitutive cooperation, given density fluctuations and positive density dependence.

### Genetic mixing can lead to coercive strategies

Next, we ask what are the effects of genetic mixing on the evolution of QS-controlled cooperation? To explore this question, we performed simulations where we varied the average number of genotype founders per local population 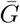, and contrasted QS-controlled and constitutive cooperation (Fig. 3A). In the clonal limit 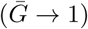, we see the efficiency benefit of matching cooperative behavior to environment compared to constitutive cooperation, as illustrated in Fig. 2A. As the degree of genotypic mixing increases, the overall payoff for constitutive cooperation is in addition fast diminishing due to the evolution of increasingly lower levels of constitutive cooperation, tending to zero. In contrast, cooperation is more robust to increased genetic mixing when controlled by QS. Note that the average payoff of QS-controlled cooperation is below the baseline, for high levels of *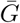*. This is because of persistent costs of low levels of signaling, maintained due to selection-drift balance.

**Fig 3.**
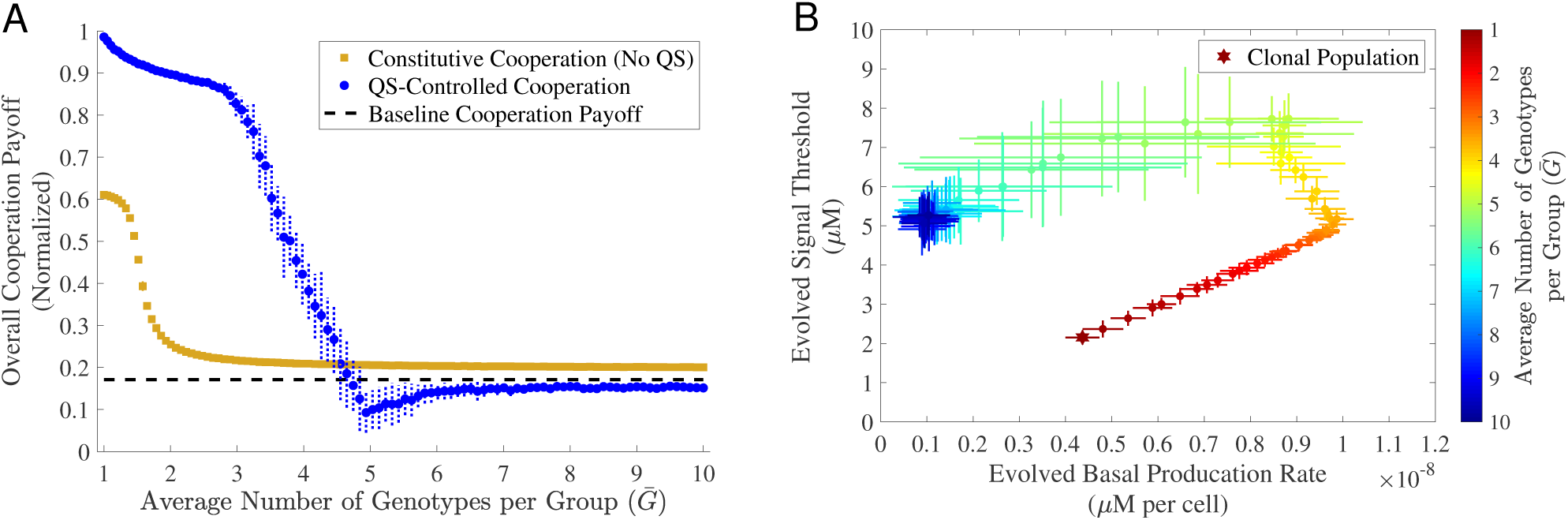
Coercive QS strategies maintain cooperation in genetically mixed populations. For both constitutive cooperation and QS-controlled cooperation, we used a fixed cost of cooperation. We also used a fixed cost of signaling in QS-controlled cooperation. In all cases, we evolved a population of 5, 000 individuals for 5, 000 generations. Each dot represents the evolved mean results (averaged over the last 50 generations) for different average number of genotypes per group 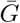. The horizontal and vertical error bars represent the standard deviation over 30 replications. **(A)** Overall cooperation payoff. The black dashed line represents the baseline payoff. **(B)** Co-evolved production rate and signal threshold for QS-controlled cooperation. The star dot represents the clonal population (*G* = 1). The remaining parameters used in the simulations can be found in *Appendix*, Table S1.

To understand the greater robustness of QS controlled cooperation we examined the joint evolution of the component signal and response traits under different conditions of genetic mixing (Fig. 3B, also see *Appendix*, Fig. S3A). Consistent with earlier theory [25], we found that clonality selected for ‘conspiratorial whisper’ strategies, coupling low signaling with low response thresholds. Conversely, under moderate genetic mixing we found the evolution of ‘coercive’ strategies featuring high signal production and high thresholds to response — capable of inducing greater cooperative responses from their less coercive ancestors when sharing a local population. For low to intermediate levels of genetic mixing 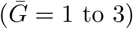, we see that the evolved genotypes stay close to the functional constraint (*pN* /*u > S_Th_*), which implies that they are able to effectively identify the density threshold when working as a solitary clone (as in Fig. 2A). However, in the event of a mixed sub-population the genotype with the higher signal and response threshold will act as a conditional cheat by inducing greater cooperative investment from its partner. As genetic mixing is further increased, the probability of ever experiencing a clonal environment is diminished (e.g. for *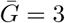*, the per sub-population probability of clonality is ∼0.17), and therefore selection on clonal efficacy is relaxed. In these low relatedness contexts, we see evolutionary trajectories towards simple cheating strategies, captured by high response thresholds and low / drifting signal production (Fig. 3B).

To explore how genetic constraints affect QS-controlled cooperation, we also investigated the overall cooperation payoff when one trait was genetically constrained to be constant. When only evolving the response threshold (holding signal production constant and non-zero), the overall cooperation payoff is rapidly lost when *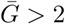* (*Appendix*, Fig. S4B). These ‘response-evolving’ populations can perform effective QS regulation in a clonal context (*G* = 1), but they are more prone to cheat takeover given genetic mixing compared to the joint-evolving populations (*Appendix*, Fig. S4A), as they cannot evolve coercive strategies. In contrast, the ‘signal-evolving’ populations (with genetically fixed response thresholds) illustrate an example of a genetic constraint driving increased cooperative robustness (*Appendix*, Fig. S4B).

### Auto-regulation sustains QS-controlled cooperation under high genetic mixing

Recent theory and experimental work has suggested that auto-regulation (specifically, a positive feedback loop between signal response and signal production [22, 47]) helps to maintain QS-controlled cooperation by increasing phenotypic assortment [32]. To investigate the role of auto-regulation in our simulation framework, we repeated the analyses of the previous section with the addition of a third evolutionary dimension, auto-regulation ratio *r* (the ratio of maximally induced production to baseline signal production (Eq. (2), see *Methods Summary*)). We first compared the overall cooperation payoff of QS-controlled cooperation with auto-regulation against QS-controlled cooperation without auto-regulation and constitutive cooperation. From Fig. 4A, we see that auto-regulation QS enhances the evolutionary robustness of cooperation in the face of medium to high levels of genetic mixing.

**Fig 4.**
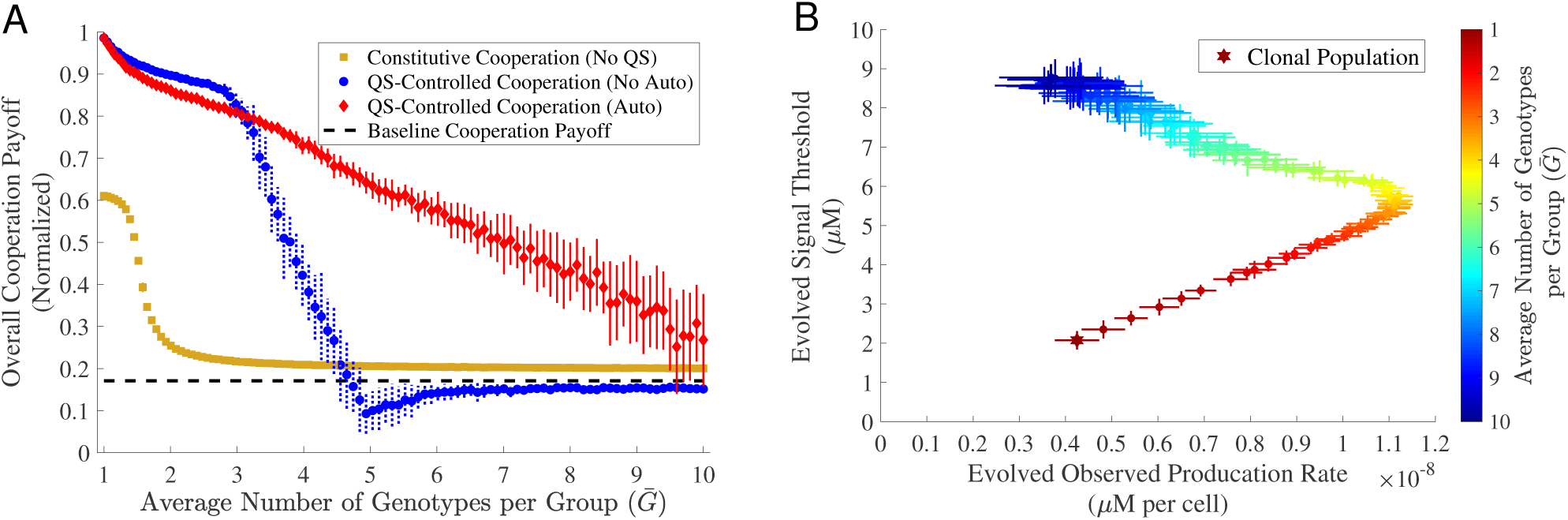
Auto-regulation extends the window of QS-controlled cooperation to greater genetic mixing. For both constitutive cooperation and QS-controlled cooperation, we used a fixed cost of cooperation. We also used a fixed cost of signaling in QS-controlled regimes. In all cases, we evolved a population of 5, 000 individuals for 5, 000 generations. Each dot represents the evolved mean results (averaged over the last 50 generations) for different average number of genotypes per group 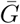. The horizontal and vertical error bars represent the standard deviation over 30 replications. (A) Overall cooperation payoff. The black dashed line represents the baseline payoff. (B) Co-evolved production rate and signal threshold for QS-controlled cooperation with auto-regulation. The star dot represents the clonal population (*G* = 1). The remaining parameters used in the simulations can be found in *Appendix*, Table S1.

To begin to decipher the mechanisms of greater resilience of QS with auto-regulation, we again examined the joint evolution of the three component traits, *p, S_Th_* and *r* (Fig. 4B, and also see *Appendix*, Fig. S3B). In *Appendix*, Fig. S5, we see that the evolved auto-regulation ratio *r* is close to 1 in the clonal context (i.e. doubling of total signal production under maximal auto-regulation, compared to baseline), suggesting there is some benefit to auto-regulation in a clonal context. In contrast, under conditions of genetic mixing auto-regulation evolves to higher levels, peaking at 8 for intermediate levels of mixing. In Fig. 4B (also *Appendix*, Fig. S5) we plot total signal production (i.e. a composite of baseline and auto-regulation behavior) against response threshold, and similarly to Fig. 3B we see the signature of a coercive escalation of signal and response threshold as genetic mixing increases. However, in contrast to Fig. 3B we now see that even at our highest levels of genetic mixing, there is still sufficient signaling and responsiveness (Fig. 4B and *Appendix*, Fig. S5) to maintain cooperative rewards above baseline (Fig. 4A).

To further challenge quorum sensing controlled cooperation, we introduced constitutive (and immutable) cheats at a certain rate in every generation (constitutive cheats have zero signal production and maximal signal threshold. See *Appendix* for more details). As can be seen from *Appendix*, Fig. S6 (also see *Appendix*, Fig. S7), the evolution of auto-regulation also protects against challenge with constitutive cheats.

### Generalized reciprocity protects QS-controlled cooperation from exploitation by cheats

To build a mechanistic understanding of why QS-controlled cooperation with auto-regulation is more robust, we measured the phenotypic assortment between individual and group cooperative investment. Fig. 5A shows that in the case of *G* = 2 (two genotypes per sub-population) and the absence of auto-induction, the relationship between the cooperative behavior of an individual (x-axis) and of its group (y-axis) is positive but with substantial variation. In contrast, the introduction of auto-induction (Fig. 5B) produces a much tighter relationship between individual and group levels of cooperation. Together, Fig. 5 (also see *Appendix*, Fig. S8) illustrates that positive auto-regulating bacteria are better able to coordinate their cooperative investment at a group level, and therefore reduce the degree of exploitative mismatches between focal individuals and other members of the group (see [32]).

**Fig 5.**
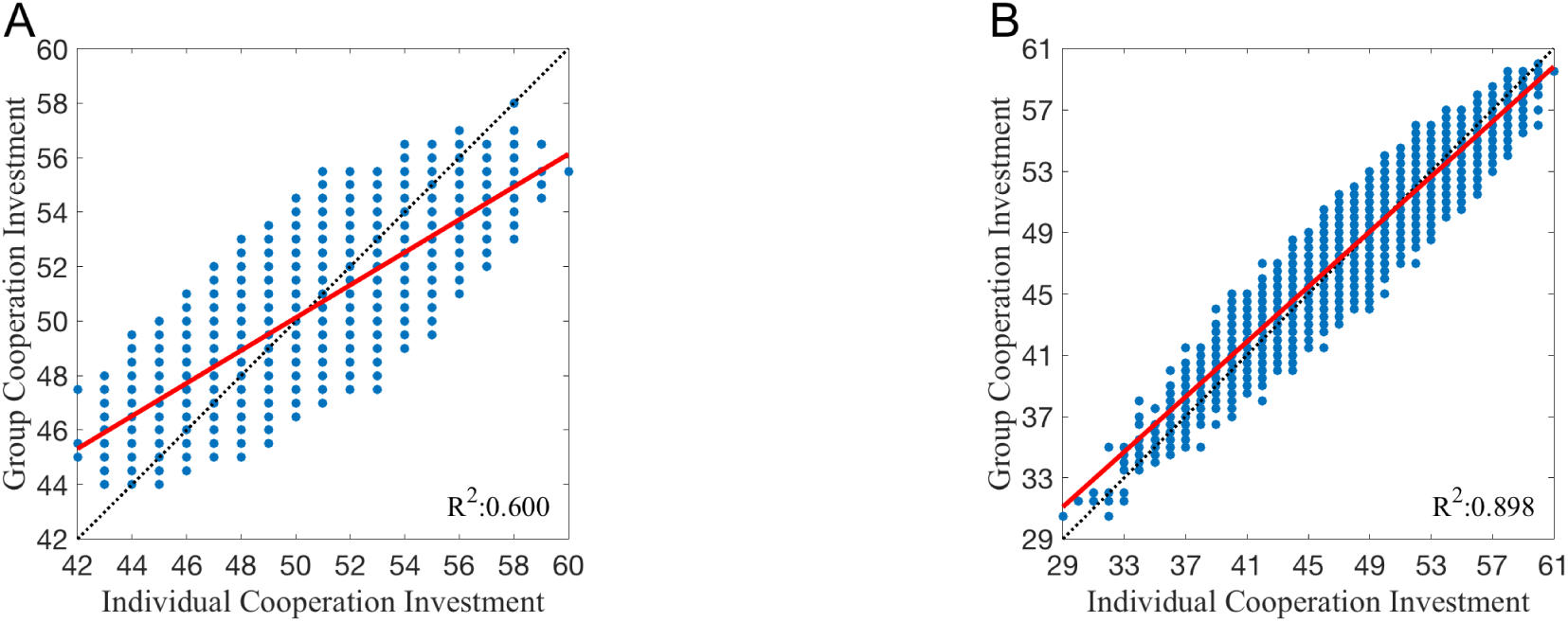
Generalized reciprocity facilitates QS-controlled cooperation. For fixed costs of cooperation and signaling with the number of mixing genotypes *G* = 2, we collected 5, 000 same initial genotypes and evolved them for 5, 000 generations with no auto-regulation (**A**) and auto-regulation (**B**). We recorded the individual and group mean investment for cooperation at the last generation over 100 replications. Each blue dot represents an individual’s investment against its group mean investment. The red lines are the regression lines fitted using the generalized linear model with a normal distribution. The analysis of covariance shows there is a significant difference between the slope of no auto-regulation in (**A**) and the slope of auto-regulation in (**B**) (*F*-test, *p* = 0.000). Similar results varying *G* can be found in *Appendix*, Fig. S8. The remaining parameters used in the simulations can be found in *Appendix*, Table S1.

To further diagnose how social selection is modified by auto-regulation we partitioned selection within and between groups (*Appendix*, Figs. S9 and S10). Consistent with Fig. 5, a levels of selection partitioning illustrates that auto-regulation acts to dramatically reduce within-group selection for cheats.

## Discussion

In this study we examined the evolutionary dynamics of quorum sensing traits in an *in silico* system, to remove the complexities of experimental model systems that have evolved under diverse and largely unknown ecological contexts. Stripping away the system specific complexities of quorum-sensing highlights that QS is at base a co-ordination game, where the reward for a particular signaling strategy depends on the prevailing strategy of response and vice versa [48, 49]. Under the defined context of our *in silico* environments, populations that exploit environments clonally can jointly tune signal and response traits to effectively resolve and respond to variation in local sub-population density, triggering cooperative investments only in sub-populations where density is above a critical threshold (Fig. 2). The introduction of genetic mixing (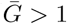 founder per sub-population) led to a broader array of strategies including ‘coercion’ (higher signal, higher response threshold, Fig. 3) and ‘generalized reciprocity’ [50] (positive auto-regulation, Fig. 4) that both independently and in conjunction contribute to the maintenance of QS regulated cooperative investments, in the face of cheats.

The description of higher signal / higher response threshold strains as ‘coercive’ is motivated by their ability to force greater degrees of cooperative response when mixed with ancestral lower response strains, while also maintaining their ability to work effectively as a clone. Kentzoglanakis *et al.* present a conceptually related *in silico* evolution model to describe a distinct biological phenomenon [51]: the evolution of plasmid intracellular copy-number control by the joint action of plasmid encoded trans-acting replication inhibitors (the signal trait), and the binding affinity of the inhibitor targets (the response trait). The authors demonstrated that the joint evolution of the signal and response trait generates increased collective efficiency (in this case, optimization of plasmid number within cells), and interpreted this model in the context of the evolutionary ‘policing’ literature [52, 53], with the repressor / signal interpreted as a ‘policing’ trait, and the target affinity / response trait interpreted as a critical and joint-evolving ‘obedience’ trait. In both the plasmid and QS contexts, we see the potential for similar co-evolutionary runaways towards increasing coercion (high signal, low response equilibria) under conditions of increased genetic mixing (Fig. 3 and also see Fig. 4 in [54]). However, as genetic mixing increases this coercive peak in signaling (policing) fails due to a collapse in obedience / response (Fig. 3). The resulting hump in signal investment with increased genetic mixing is predicted by a simple analytical game theory model [25] and now has support from two distinct simulation models built with very different biological motivations (this study and [51, 54]), which raises the challenge of why this result has been difficult to pin down experimentally, despite explicit attention [55]. Later in the discussion we return to this point in a general overview of the empirical context, but in short, it appears that the genetics of auto-regulation present an effective mechanistic block to the elaboration of coercive strategies.

One of the key hallmarks of many (but not all) QS regulatory architectures is signal auto-regulation, where signal response is coupled to increase the signal production [40, 56–59], leading to increased synchrony across individual cellular responses [47]. To explore the evolutionary role of auto-regulation in our system, we added auto-regulation as a third evolving trait, and found that this additional evolutionary dimension led to a further increase in the robustness of QS controlled cooperation (Fig. 4). In the evolved auto-regulation lineages we found a stronger degree of phenotype matching (assortment) between individuals and their group (Fig. 5), demonstrating that positive feedback control of signal production allows individuals to tune their cooperative behavior to their social environment. This result is consistent with a recent experimental paper on *P. aeruginosa*, which demonstrated that *P. aeruginosa* can facultatively tune its per-capita cooperative investment to the proportion of wildtype cooperators in its local group, in a manner that will promote the maintenance of cooperation [32]. Allen *et al.* described this behavior as an example of generalized reciprocity, highlighting that by encoding a simple rule of ‘cooperate when with cooperators’ bacteria can increase the robustness of cooperation and the regulatory architectures that control cooperation [32].

In the simple environmental and genetic world of our *in silico* bacteria, populations readily evolve complex strategies of coercion and generalized reciprocity. While generalized reciprocity has been reported for *P. aeruginosa*, coercion has been far more elusive, despite direct experimental evolution tests [55]. Popat *et al.* experimentally evolved *P. aeruginosa* under conditions of high and low genetic mixing, and found that under conditions of intermediate and low genetic mixing, the level of signal production only went down (alongside response); there was no peak in coercion [55]. One possible account for this disconnect with our simulations is that on the ∼ 1 month timescale of experimental evolution the evolutionary dynamics are constrained by the genetic mechanisms of auto-regulation: The easiest solution to reduce signal response is to mutate the signal receptor (in *P. aeruginosa*, this is frequently achieved by Δ*lasR* mutations) which has the pleiotropic consequence of also largely abolishing signal production.

This argument suggests that coercive strategies are more likely to be evolvable on short timescales in bacteria without strong auto-regulatory constraints, such as *V. cholerae* [60] (but see [61]). In our main text results all traits could independently evolve, and thus both generalized reciprocity (signal auto-regulation) and coercion (high signal / low response) are accessible simultaneously. In *Appendix*, Fig. S4, we introduced simple genetic constraints (constraining the evolution of one trait and allowing others to freely evolve) and found substantial shifts in evolutionary trajectories, either helping (with a fixed response, see blue dots in *Appendix*, Fig. S4B) or harming (with a fixed signal, see yellow dots in *Appendix*, Fig. S4B) the maintenance of cooperation depending on genetic details.

The existence of a genetic constraint does not imply that over longer time-scales the constraint is immutable. Take for example the constraint imposed by *lasR* co-regulation on the trajectories of signal production and signal threshold in *P. aeruginosa*. Sandoz *et al.* reported two *lasR* mutants (*lasR*5 and *lasR*8) that displayed near-wildtype levels of signal production but with lower level of signal response [27]. In principle, it is possible that signal production and response could evolve independently in *P. aeruginosa* by separately targeting steps that are downstream of *lasR*, for instance targeting multiple promoter sites to separately tune the impact of LasR on signal synthase and cooperative effector genes. Gurney *et al.* [33] recently demonstrated using experimental evolution that *P. aeruginosa* can rewire its response to multiple signal inputs in order to escape ancestral genetic constraints on social behaviors — in this example, through the evolution of novel cheating strategies to escape pleiotropic constraints termed ‘metabolic incentives to cooperate’ [29].

In our ‘*in silico*’ evolution, we know the ecological challenges that bacteria are facing in the controlled environments. Specifically, we defined a density threshold for the rewards of turning on cooperation and showed that bacteria can evolve strategies that are adaptations to ‘density sensing’. However, as a result we inevitably also evolve spandrels (a byproduct of adaptive selection, see [62]). For example, our evolved bacteria can in principle perform a ‘diffusion sensing’ role to differentiate mass transfer regimes [20], despite never experiencing this challenge. On the other hand, it is possible that if we set the environmental challenges to ‘diffusion sensing’, we will evolve ‘density sensing’ as spandrels (or exaptation). The ability to precisely define and control the environment of adaptation and the genetic constraints of the ancestor suggest that *in silico* bacteria are a fruitful model for the study of adaptation and exaptation in quorum-sensing bacteria.

## Supporting information

**S1 Appendix.** Supplementary information.

## Acknowledgments

This work was supported by Simons Foundation Grant 396001 (to S.P.B.).

## Appendix

### Simulation model

In the *in silico* evolution, we consider two evolving traits, basal production rate (*p*) and signal response threshold (*S_Th_*) for simulations in the absence of auto-regulation, whereas we introduce an additional evolving trait, auto-regulation ratio (*r*) for simulations including the auto-regulation mechanism. Each individual in the population pool has a single genotype which consists of those two or three evolving traits. The individuals can make their own decisions to turn on or off cooperation as a function of signal mediated interactions, which in turn depend on the physical and social environment. Specifically, the evolution process is described in *Main Text*, Fig. 1:

1. Total *N_pop_* genotypes with same initial conditions (same 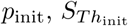 and *r*_init_) were generated to from a population pool.
2. A certain number of genotypes (*G*, drawn from zero-truncated Poisson distribution, unless otherwise specified) were randomly selected (with replacement) from the population pool and form a mixed sub-population.
3. For each of *N_env_* sub-population testing environments, the signal concentration in the mixed sub-population can be calculated as *S*^*^ using Eq. (S2) (or Eq. (S7) for auto-regulation case).
4. Each genotype in the mixed sub-population was evaluated for its overall cooperation payoff separately across all sub-population testing environments (where the cellular density was varied) using Eq. (S3) (or Eq. (S8) in auto-regulation case): Each individual paid for its own cost for signaling and the cost of cooperation, if any, but only gained a benefit when the number of cooperators in sub-population were greater than a certain threshold, *N_Th_*.
5. Repeat 2) to 4) until the same size of population pool was formed.
6. All individuals were selected (with replacement) from the population pool to reproduce with a probability proportional to their overall cooperation payoff.
7. All evolving traits (*p, S_Th_* and *r*) of the offspring were subject to mutation at rates *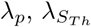* and *λ_r_* with standard deviations *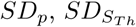* or *SD_r_* for different traits. Specifically, for each evolving trait, the actual number of individuals selected for mutation was drawn from a Poisson distribution with the mean being *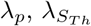* and *λ_r_*, respectively. The mutation operation was done by adding a value of *N* (0, *SD*) to the original trait value, where *N* (0, *SD*) is the normal distribution with a mean 0 and a standard deviation *SD* to be *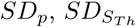* or *SD_r_* for different traits.
8. Repeat 2) to 7) until *Gen*_max_ generations were reached.

### Model assumptions

For the computational models presented in the paper, we assume that:

1. A single signal type exists.
2. The environment in each sub-population forms a closed, i.e., no mass transfer.
3. For a given sub-population testing environment, an individual can be rewarded with a benefit for cooperation only if the number of cooperators is greater than the defined threshold (*N_Th_*).
4. All offspring cloned from a certain parental genotype behave similarly, i.e., no heterogeneity.
5. The signal concentration in each sub-population rapidly reaches equilibrium estimated by Eq. (S1) or Eq. (S6), depending on whether invoking the auto-regulation mechanism.

### Computational model of quorum sensing without auto-regulation

The computational model of signal dynamics for quorum sensing without auto-regulation is given as below:

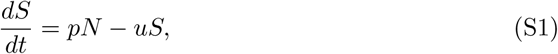

where *S* is the local signal concentration, *t* is time, *N* is the local cell density, *p* is the basal signal production rate, and *u* is the signal decay rate. The equilibrium of Eq. (S1) is given by:

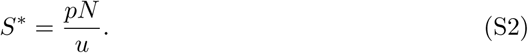

In the absence of the auto-regulation mechanism, the individual genotype’s overall cooperation payoff across all sub-population testing environments is assessed as follows:

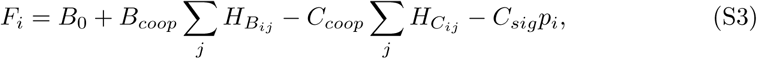

where *i* (*i* = 1, 2, *…, N_pop_*) represents an individual genotype, *j* (*j* = 1, 2, *…, N_env_*) represents the index number of a sub-population testing environment, *B*_0_ is the baseline payoff, *B_coop_, C_coop_, C_sig_* are constants for the benefit of cooperation, cost of cooperation and cost of signaling, respectively, *p_i_* is the basal signal production rate of the genotype *i*. The function of cooperation cost of the individual *i* in the sub-population testing environment *j* is defined as:

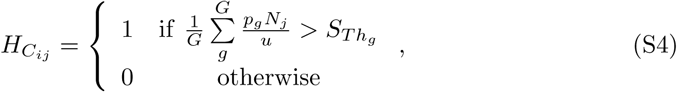

where *G* is the number of mixing genotypes in a sub-population, *N_j_* is the local cellular density in the *j*^*th*^ sub-population testing environment, and *p_g_* and *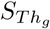* are the signal production rate and signal response threshold of the genotype *g* (*g* = 1, 2,…, *G*) in the sub-population, respectively. Similarly, the function of cooperative benefit of the individual *i* in the sub-population testing environment *j* is defined as:

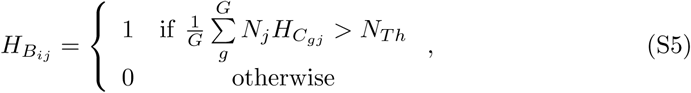

where *N_Th_* is the cellular density threshold (defined as the median cellular density across all testing environments).

### Computational model of quorum sensing with auto-regulation

The computational model of signal dynamics for quorum sensing with auto-regulation is given as below:

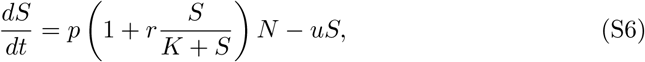

where *S* is the local signal concentration, *t* is time, *N* is the local cell density, *p* is the basal signal production rate, *r* is the ratio of auto-regulation production to basal signal production, *K* is the half concentration value, and *u* is the signal decay rate. The equilibrium of Eq. (S6) is given by:

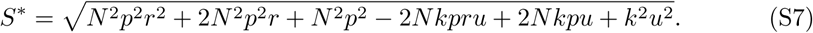

Note that previous studies have indicated the choice of Hill function exponent to be 2 [1, 2]. However, for the purpose of computational convenience, we used 1 as the Hill function exponent, which can lead to a close form solution, Eq. (S7). When invoking the auto-regulation mechanism, the individual genotype’s overall cooperation payoff across all testing environments is assessed as follows:

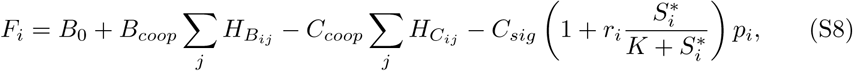

where *i* (*i* = 1, 2, *…, N_pop_*) represents an individual genotype, *j* (*j* = 1, 2, *…, N_env_*) represents the index number of a sub-population testing environment, *B*_0_ is the baseline payoff, *B_coop_, C_coop_, C_sig_* are constants for the benefit of cooperation, cost of cooperation and cost of signaling, respectively, *p_i_* and *r_i_* are the basal signal production rate and auto-regulation ratio of the genotype *i*, respectively, and *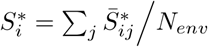* where *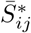* is defined in Eq. (S9). The function of cooperation cost of the individual *i* in the sub-population testing environment *j* is defined as:

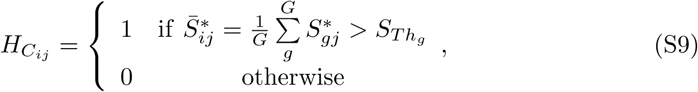

where *G* is the number of mixing genotypes in a sub-population, *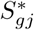* (calculated by Eq. (S7)) and *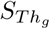* are the equilibrium signal concentration and signal response threshold of the genotype *g* (*g* = 1, 2,…, *G*) in the sub-population, respectively. The function of cooperative benefit of the individual *i* in the sub-population testing environment *j*, and *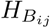* is defined as the same as in Eq. (S5).

### Adding noise to signal

To investigate how clonal populations cope with signal noise to sustain cooperation, we added noise to the equilibrium signal. In the simulations, the noise signal is drawn from a normal distribution with mean *S*^*^ and standard deviation *κ S\**, i.e., *N* (*S*, κ S\**), where *S\** is the equilibrium signal calculated from Eq. (S2) or Eq. (S7) and *κ* is a constant indicating the strength of noise. Note we set all negative values for the noise signal to be 0.

### Generating genetic mixing

In the evolution simulations where we varied the genetic relatedness, different numbers of mixing genotypes were introduced to form sub-populations to evaluate the overall cooperation payoff for each genotype in every generation. Unless specified otherwise, the actual number of mixing genotypes in each sub-population in every generation was drawn from a zero-truncated Poisson distribution with the average being 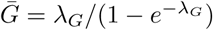, where (*λ_G_ ϵ* [0, 10]; step size: 0.1). We define the clonal case (*G* = 1) when *λ_G_* = 0 where exact one genotype will be selected to form the sub-population, i.e., no genetic mixing. Note that the number of sub-populations may be different in every generation due to the variation of mixing genotypes in each subpopulation.

### Constructing constitutive cooperators

To investigate how decision making interact with social behaviors of cooperation, we compared the overall payoff of cooperation of individuals mediated by QS with those in the absence of collective control. Specifically, we constructed constitutive cooperators which do not have the ability to make social informed choices. In the clonal case, wild-type individuals will always cooperate. This will incur a penalty to each of such individuals for cooperating in wrong environments^1^. In the genetic mixing scenarios, the cooperative benefits of wild-type individuals will be shared evenly with all group members. In the simulations, all individuals were subject to mutation, switching from a wild-type to mutant, or mutant to wild-type depending on their own initial type. The actual number of replacement individuals was drawn from a Poisson distribution with the mean being 0.01. Note that mutant individuals will always reap the benefits of cooperation without paying for any cost. Formally, the overall cooperation payoff in the constitutive cooperation scenarios can be defined as:

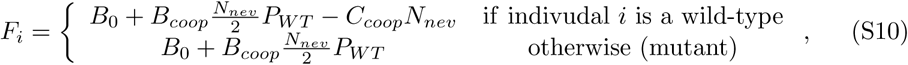

where *P_W T_* is the proportion of wild-type individuals in the sub-population with a group size *G*.

### Introducing constitutive cheats

To challenge the quorum sensing system, in the simulations of invasion by a cheat phenotype, we replaced a certain number of individuals^2^ chosen at random in the population pool with constitutive cheats in every generation at a certain rate. The constitutive cheat is defined as a genotype with a zero basal production rate and a maximum possible signal threshold. In other words, a constitutive cheat does not produce or respond to signal. The actual number of cheats introduced into each generation is drawn from a Poisson distribution with the mean being *λ_Cheat_*. Note that the constitutive cheats are both immutable, which means they cannot be eliminated through mutation, and inheritable, which means their offspring are still cheats.

### Measuring phenotypic assortment of cooperative investment

To test if the auto-regulation mechanism could be explained by the generalized reciprocity theory, we recoded the mean value of cooperative investment^3^ within each sub-population in the genetic mixing scenario where individuals are grouped into small collectives. We then plotted the group mean cooperative investment against individual cooperative investment. Finally, the regression line was fitted using the generalized linear model with a normal distribution. The slope of the regression line indicates the phenotypic assortment of cooperative investment. When the slope is high, the behaviors among individuals shifts closer to each other, investing more in cooperation. Otherwise, the behaviors of investment for cooperation vary among individuals.

### Partitioning selection on cooperative investment

To further uncover the influence of the auto-regulation mechanism on cooperative behaviors in our evolution simulations, we employed the powerful conceptual framework of the Price equation to partition the selection on cooperative investment into both individual (within sub-populations) and group (between sub-populations) level [3]. The Price equation describes the change in the average amount of a trait (*z*) from one generation to the next (Δ*z*) as a function of the covariance of between the fitness and the trait value among individuals (Cov(*w_i_, z_i_*)), and the expected change in the amount of the trait value (E(*w_i_*Δ*z_i_*)) due to transmission error such as genetic drift, mutation bias, etc. The general form of the Price equation is given as below

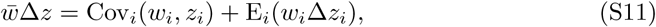

where *z* represents the trait cooperative investment, *w_i_* is the number of offspring (fitness) produced by the individual 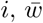 is the mean number of offspring produced, Δ*z_i_* represents the difference between the average *z* value among the individual *i*’s offspring and *i*’s own *z* value, Cov_*i*_(·) and E_*i*_(·) denote the expectation and covariance over all individuals *i* in the population respectively.

By introducing the genetic mixing in the simulations, individuals in the population have been assigned into small groups. We are able to further partition that selection based on cooperative investment to account for individuals that are nested within collectives. Specifically, we can expand Eq. (S11) by substituting its right hand side of the equation into the expectation term. Note that the groups that form each subpopulation *g* are the individuals *ig*. We can re-write the two-level Price equation as follows:

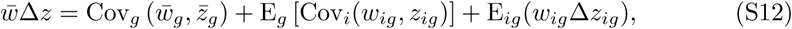

where 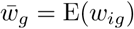. and 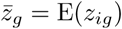 The first covariance term on the right hand side of the equation indicates the selection on cooperative investment at level of subpopulations (between-group selection), whereas the second expectation term captures the selection at individual level (within-group selection).

---

**Fig S1.**
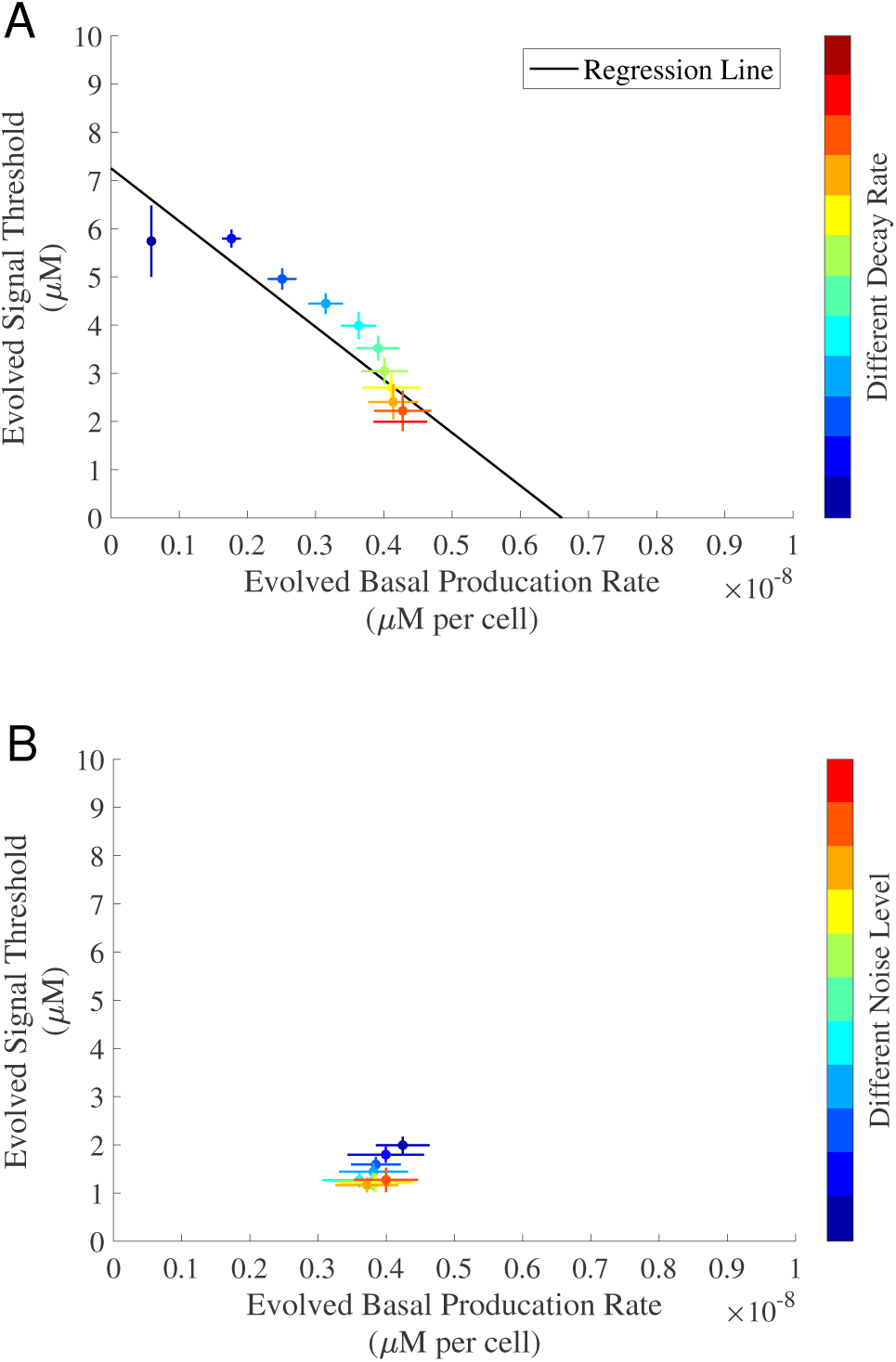
The evolved traits against signaling cost, decay rate and noise for QS-controlled cooperation. We evolved 5, 000 initially identical genotypes for 5, 000 generations. In all simulations, the cost of cooperation was fixed and there was no auto-regulation. Specifically, the population was evolved under three regimes: **(A)** Varying a range of decay rates (*uϵ* [5 × 10^−6^, 1.15 × 10^−4^]; step size: 10^−5^) with a fixed signaling cost and no noise (*κ* = 0), and **(B)** Varying levels of noise (*k* ∊ [0, 1]; step size: 0.1) with a fixed signaling cost and a fixed decay rate (*u* = 1.05 × 10^−4^). Each dot represents the evolved mean results (averaged over the last 50 generations). The color-bars indicate different values of *u* and *κ* from low (dark blue) to high (dark red) in **(A)** and **(B)**, respectively. The solid black line in **(A)** is the regression line fitted using the generalized linear model with a normal distribution: *R*^2^ = 0.824, *F*-test, *p* = 4.443 × 10^−5^. The horizontal and vertical error bars represent the standard deviation over 30 replications. The remaining parameters used in the simulations can be found in Table S1.

**Fig S2.**
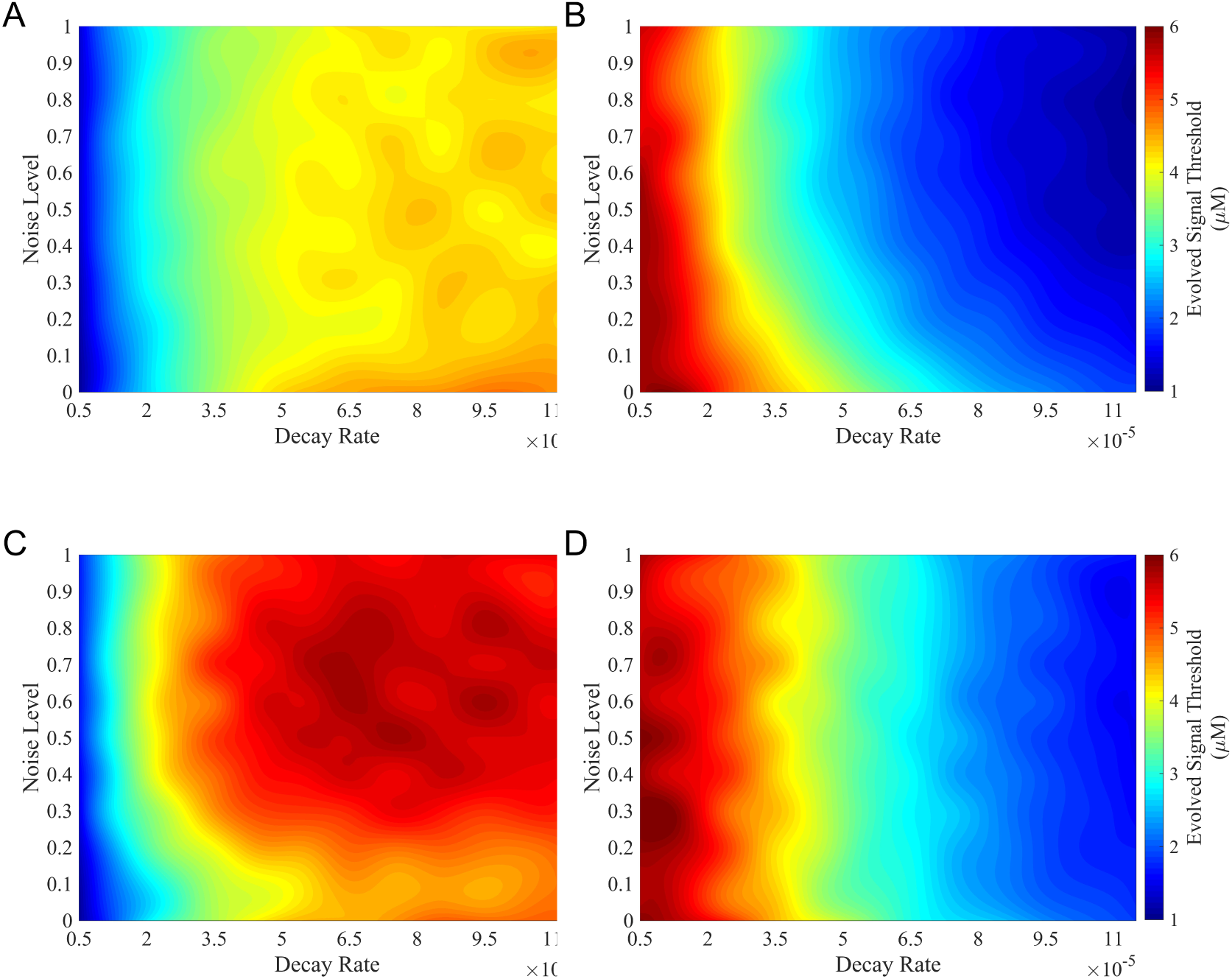
The evolved traits under different decay rates and levels of noise. We evolved 5, 000 initially identical genotypes for 5, 000 generations. In all simulations, the cost of cooperation and the cost of signaling were fixed. The decay rates, *u*, were varied in [5 × 10^−6^, 1.15 × 10^−4^] (step size: 10^−5^), and the levels of noise, *κ*, were varied in [0, 1] (step size: 0.1). The results of basal and observed production rates were reported in **(A)** and **(C)** for QS with no auto-regulation and auto-regulation, respectively. The results of signal threshold were reported in **(B)** and **(D)** for QS with no auto-regulation and auto-regulation, respectively. All reported results were averaged over 30 replications. Note that surfaces were smoothed using the spline interpolation method. The remaining parameters used in the simulations can be found in Table S1.

**Fig S3.**
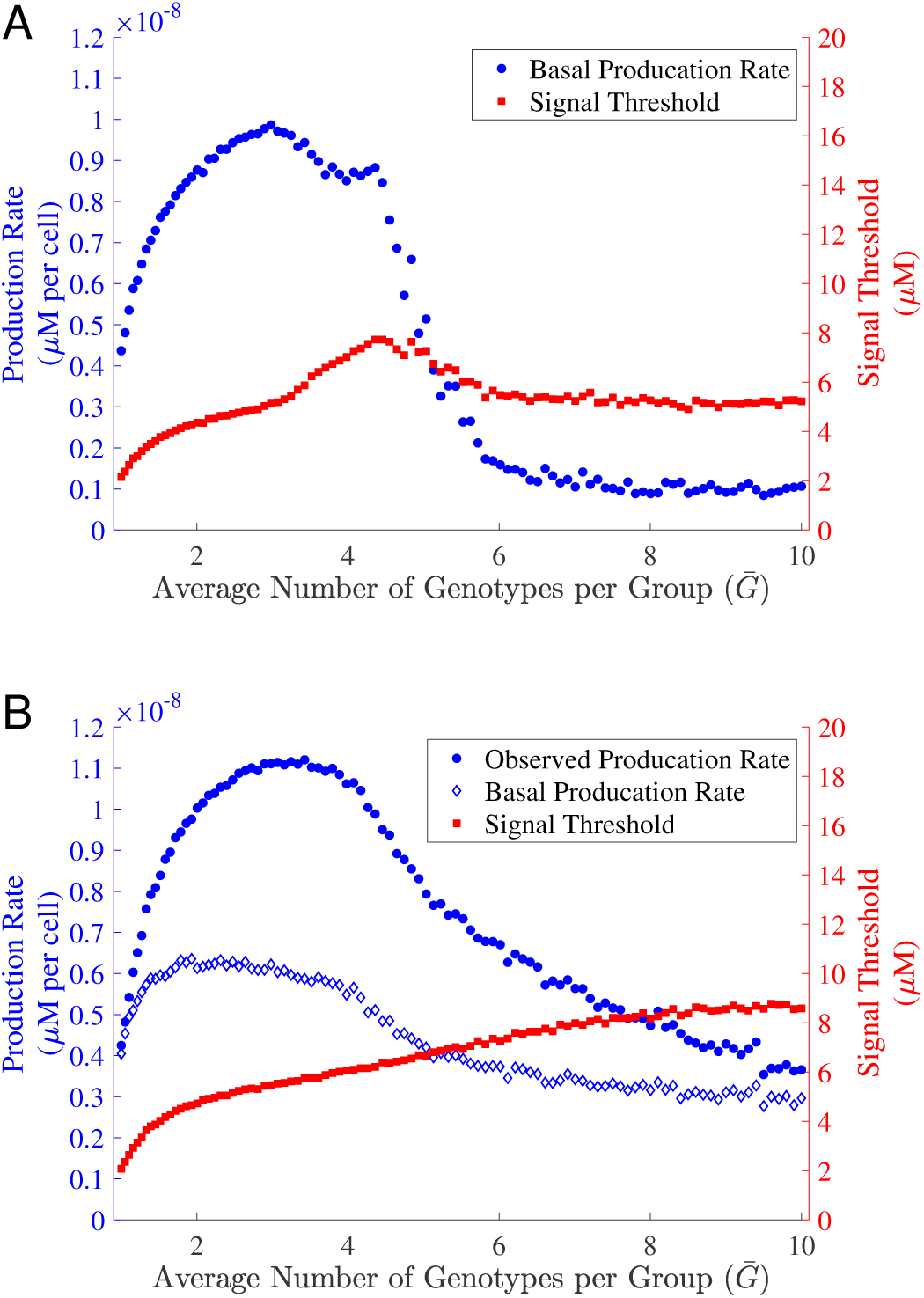
Evolved traits against genetic relatedness. We evolved 5, 000 initially identical genotypes for 5, 000 generations with no auto-regulation **(A)** and auto-regulation **(B)**, respectively. In all simulations, the cost of cooperation and the cost of signaling were fixed. Each dot represents the evolved mean results (averaged over the last 50 generations) for different average number of genotypes per group 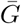 (*λ_G_ ϵ* [0, 10]; step size: 0.1). The remaining parameters used in the simulations can be found in Table S1.

**Fig S4.**
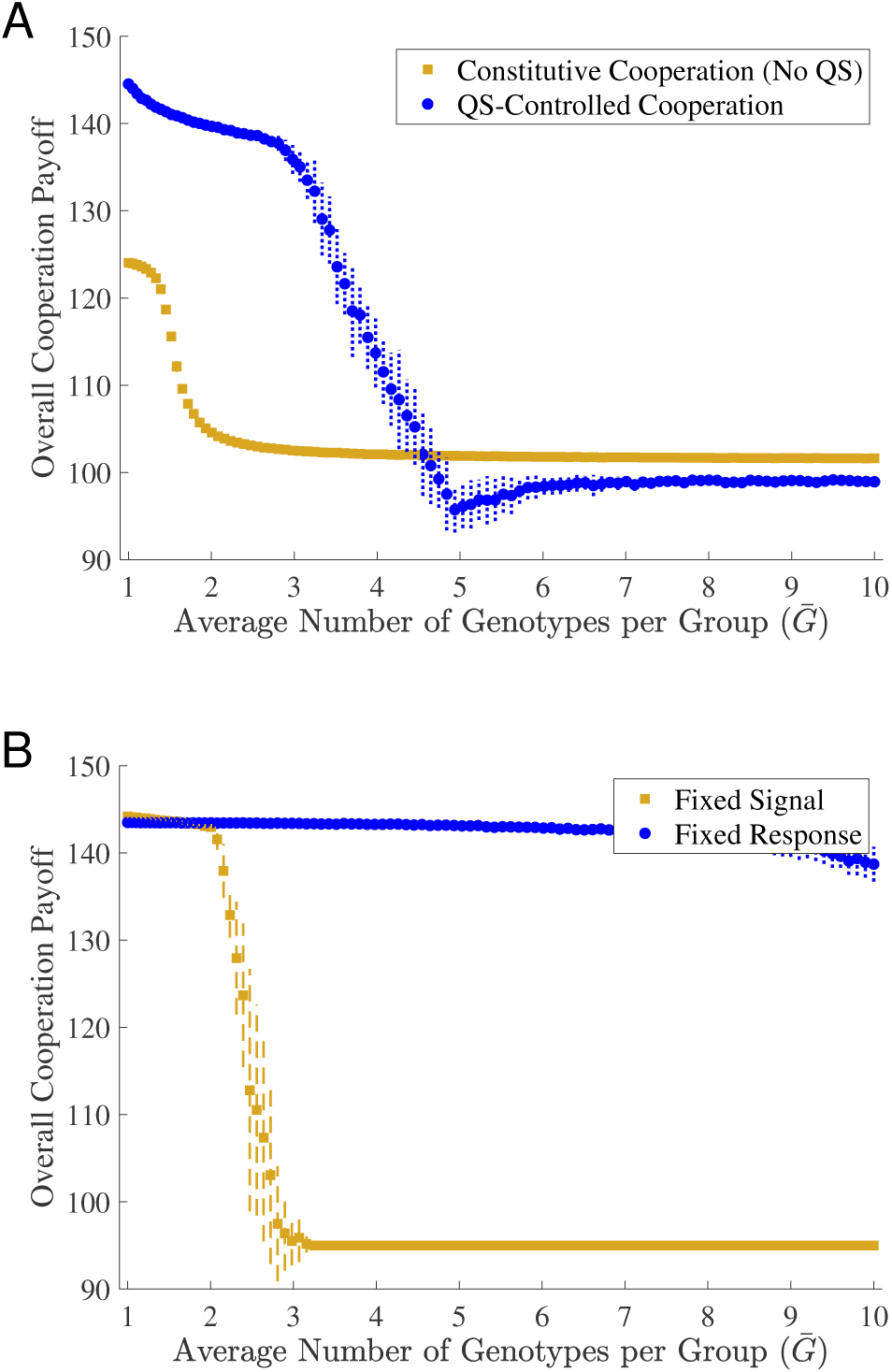
Comparison of overall cooperation payoff in different evolution scenarios. **(A)** Overall cooperation payoff of constitutive cooperation and QS-controlled cooperation. Note this figure is the same one as in *Main Text* Fig. 3A, except we used the original values of the overall cooperation payoff. **(B)** Overall cooperation payoff for fixed trait evolution. We used a fixed signal production rate of 0.5 × 10^−8^ and a fixed response threshold of 3 *μ*M for QS-controlled cooperation without auto-regulation, respectively. The cost of cooperation and cost of signaling were also set to be the same as in **(A)**. In all cases, we evolved 5, 000 initially identical genotypes for 5, 000 generations. Each dot represents the evolved mean results (averaged over the last 50 generations) for different average number of genotypes per group 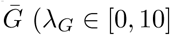; step size: 0.1). The vertical error bars represent the standard deviation of overall cooperation payoff over 30 replications. The remaining parameters used in the simulations can be found in Table S1.

**Fig S5.**
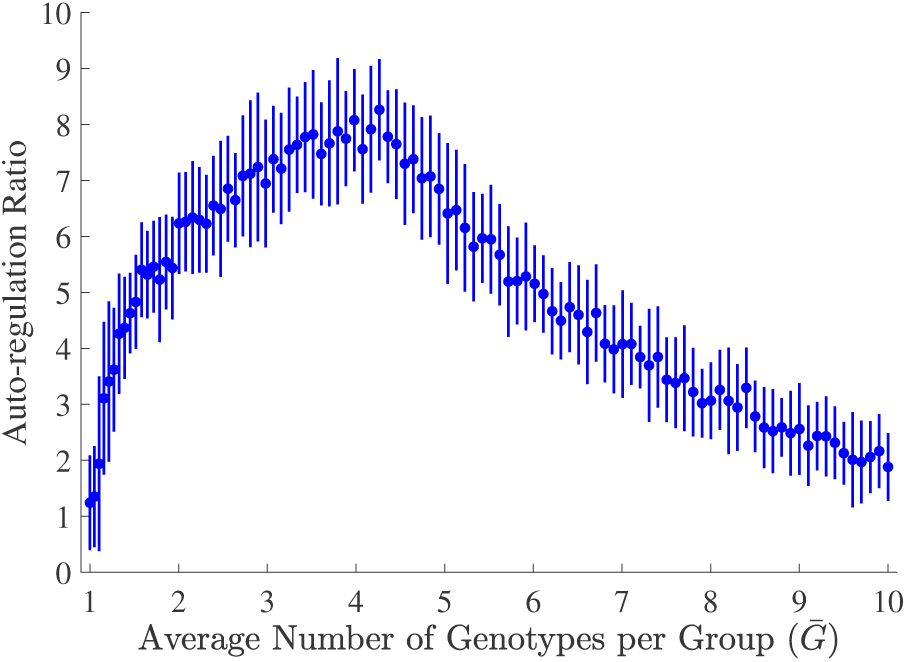
Evolved auto-regulation ratio against genetic relatedness. We evolved 5, 000 initially identical genotypes for 5, 000 generations with auto-regulation. In the simulations, the cost of cooperation and the cost of signaling were fixed. Each dot represents the evolved mean results (averaged over the last 50 generations) of auto-regulation ratio (*r* as in Eq. (S6)) for different average number of genotypes per group [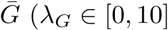]; step size: 0.1). The vertical error bars represent the standard deviation over 30 replications. The remaining parameters used in the simulations can be found in Table S1.

**Fig S6.**
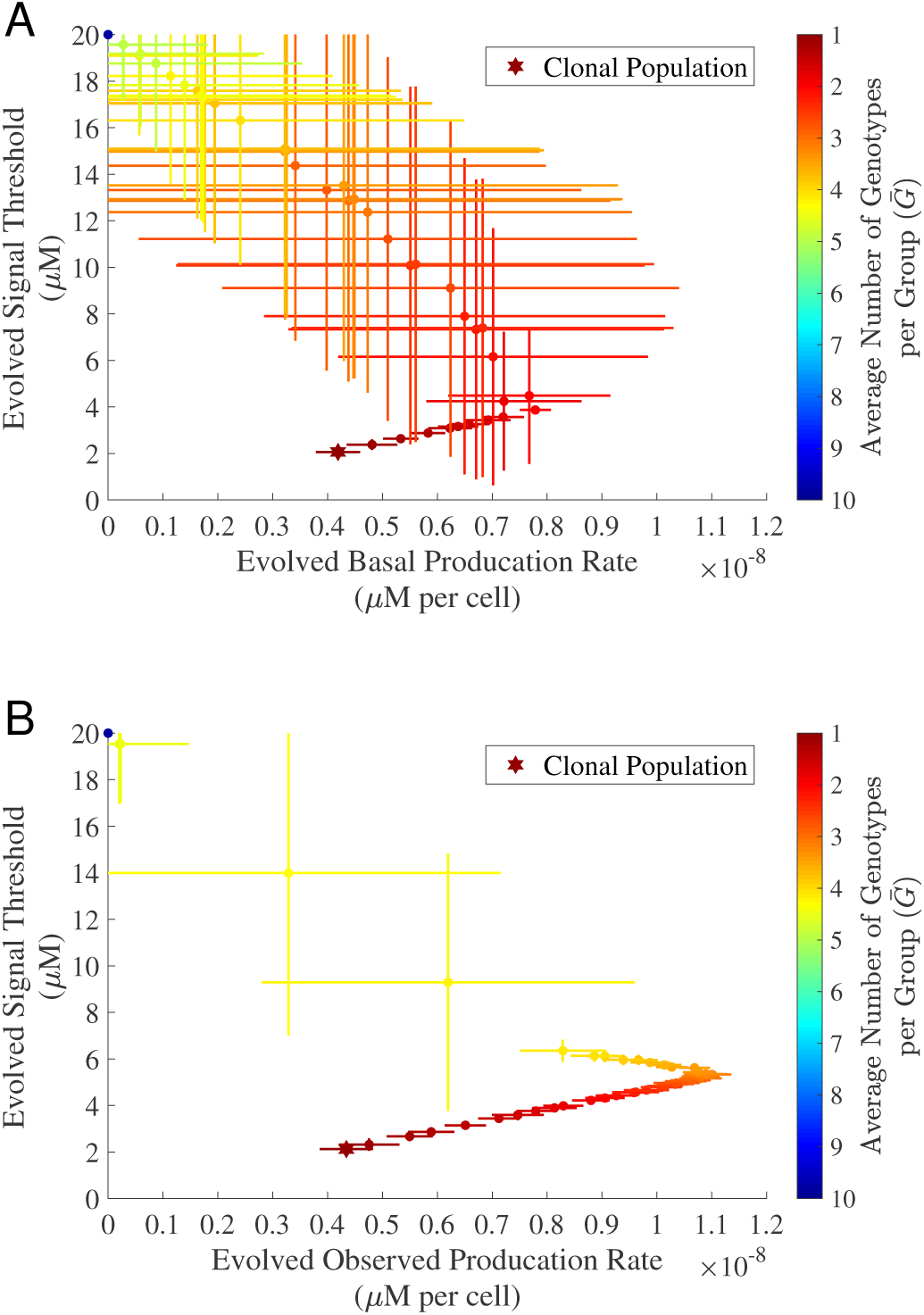
Invasion of constitutive cheats to the system with quorum sensing. We evolved 5, 000 initially identical genotypes for 5, 000 generations with no auto-regulation **(A)** and auto-regulation **(B)**, respectively. In all simulations, the cost of cooperation and the cost of signaling were fixed. A certain number of individuals (drawn from a Poisson distribution with *λ_Cheat_* = 0.1) chosen at random were replaced with the constitutive cheats in every generation. Each dot represents the evolved mean results (averaged over the last 50 generations) for different average number of genotypes per group 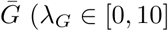; step size: 0.1), indicated in the color-bar on the right. The star dot represents the clonal case when *G* = 1. The horizontal and vertical error bars on each dot represent the standard deviation of the results over 30 replications. The remaining parameters used in the simulations can be found in Table S1.

**Fig S7.**
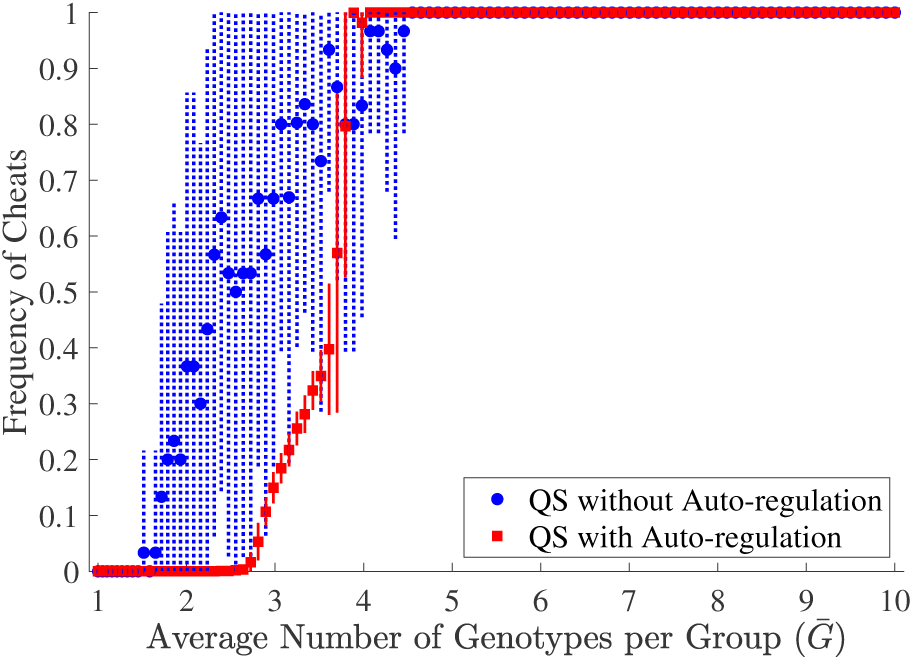
Comparison of frequency of cheats for the evolved system with or without auto-regulation. We evolved 5, 000 initially identical genotypes for 5, 000 generations with no auto-regulation **(A)** and auto-regulation **(B)**, respectively. In all simulations, the cost of cooperation and the cost of signaling were fixed. A certain number of individuals (drawn from a Poisson distribution with *λ_Cheat_* = 0.1) chosen at random were replaced with the constitutive cheats in every generation. Each round dot (no auto-regulation) or square dot (auto-regulation) represents the evolved mean results (averaged over the last 50 generations) for different average number of genotypes per group 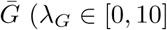; step size: 0.1). The vertical error bars represent the standard deviation over 30 replications. The remaining parameters used in the simulations can be found in Table S1.

**Fig S8.**
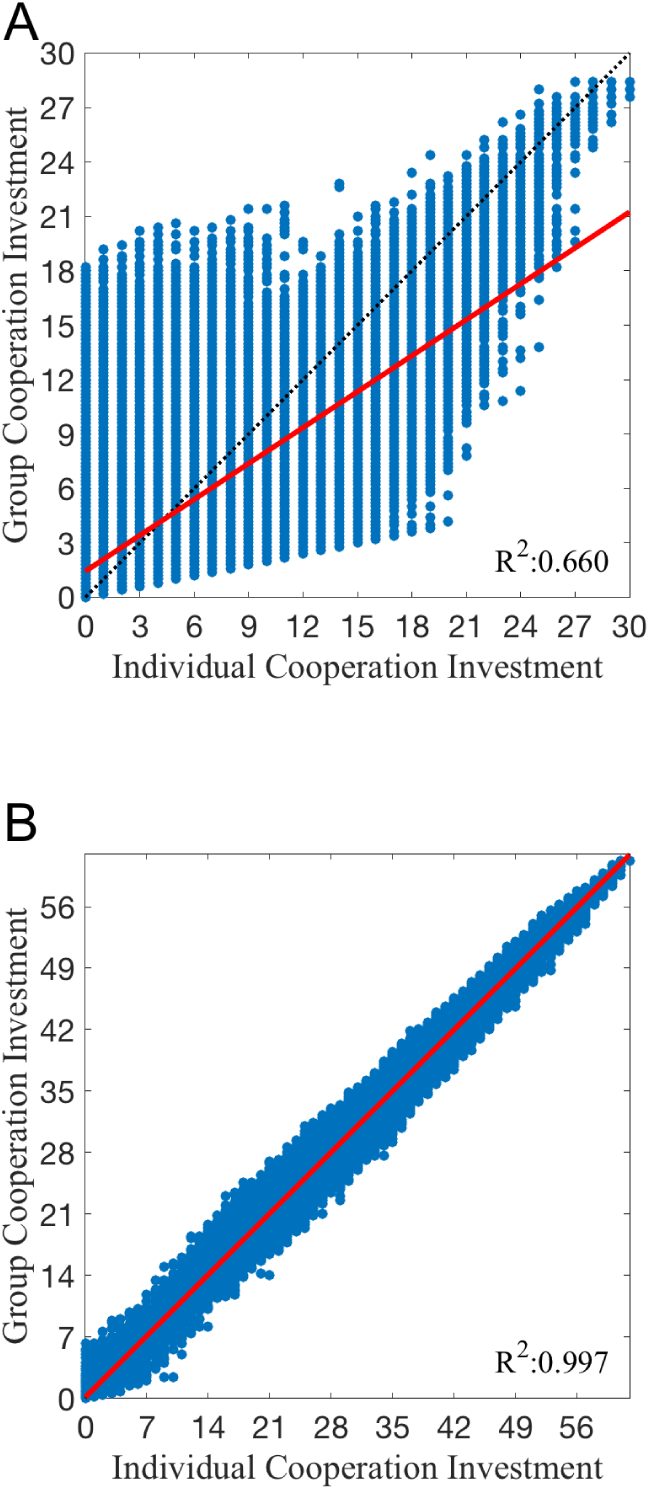
Regression analysis for individual and group mean investment for cooperation (*G* = 5). For fixed costs of cooperation and signaling with the number of mixing genotypes *G* = 5, we collected 5, 000 same initial genotypes and evolved them for 5, 000 generations with no auto-regulation **(A)** and auto-regulation **(B)**. We recorded the individual and group mean investment for cooperation at the last generation over 100 replications. Each blue dot represents an individual’s investment against its group mean investment. The red lines are the regression lines fitted using the generalized linear model with a normal distribution. The analysis of covariance shows there is a significant difference between the slope of no auto-regulation in **(A)** and the slope of auto-regulation in **(B)** (*F*-test, *p* = 0.000). The remaining parameters used in the simulations can be found in Table S1.

**Fig S9.**
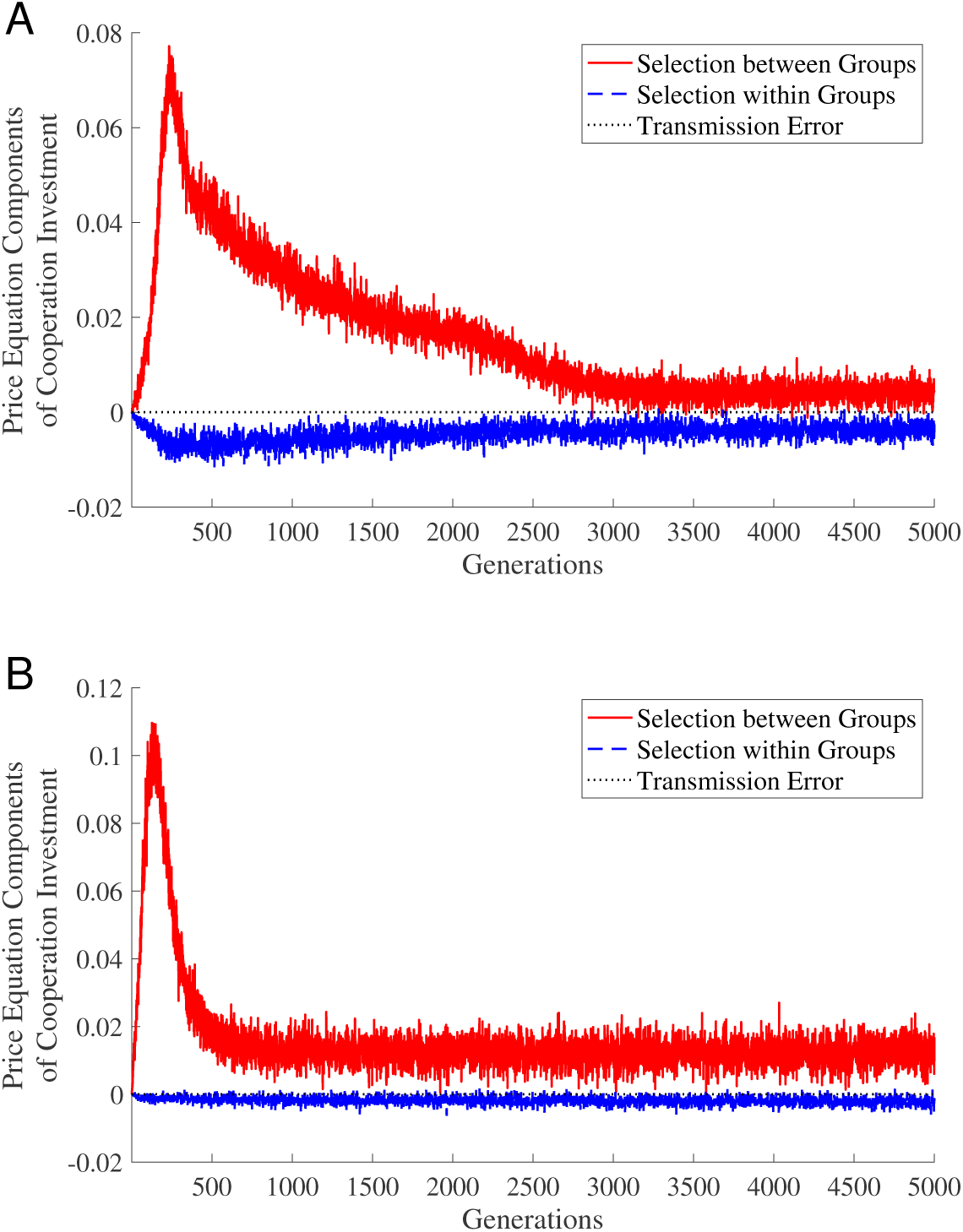
Selection on cooperative investment within and between groups (*G* = 2). We evolved 5, 000 initially identical genotypes for 5, 000 generations with no auto-regulation **(A)** and auto-regulation **(B)**, respectively. In all simulations, the cost of cooperation and the cost of signaling were fixed, and the number of mixing genotypes was fixed *G* = 2. We recorded the two-level Price equation components in every generation. The reported results were the average value over 100 replications. The remaining parameters used in the simulations can be found in Table S1.

**Fig S10.**
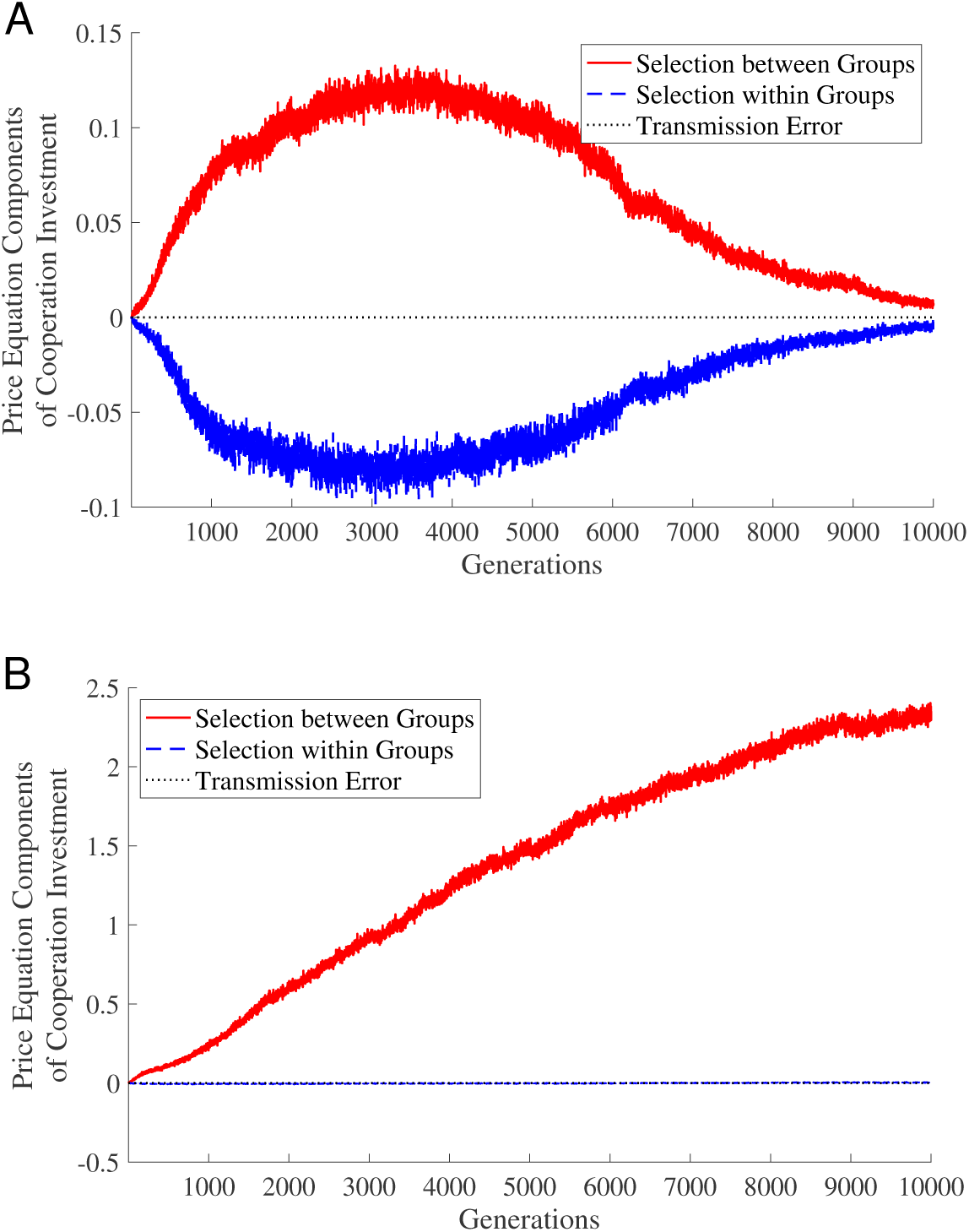
Selection on cooperative investment within and between groups (*G* = 5). We evolved 5, 000 initially identical genotypes for 5, 000 generations with no auto-regulation **(A)** and auto-regulation **(B)**, respectively. In all simulations, the cost of cooperation and the cost of signaling were fixed, and the number of mixing genotypes was fixed *G* = 5. We recorded the two-level Price equation components in every generation. The reported results were the average value over 100 replications. The remaining parameters used in the simulations can be found in Table S1.

**Table S1.** List of Model Parameters

| Symbol | Description | Default Value |
| --- | --- | --- |
| $B_0$ | baseline payoff | 100 FU<sup>1</sup> |
| $B_{coop}$ | constant value for the benefit of cooperation | 1.5 FU per CTE<sup>2</sup> |
| $C_{coop}$ | constant value for the cost of cooperation | 0.5 FU per CB<sup>3</sup> |
| $C_{sig}$ | constant value for the cost of signaling | 10<sup>9</sup> FU per $\mu\text{M}$ |
| $Gen_{\max}$ | the maximum of generations | 5000 |
| $K$ | half concentration value | 50 $\mu\text{M}$ |
| $G$ | number of mixing genotypes, unless otherwise specified, drawn from<br>a zero-truncated Poisson distribution with the average being $\bar{G} = \lambda_G / (1 - e^{-\lambda_G})$ | 1 |
| $N_{Th}$ | threshold of cellular density | 50016 cells per $\mu\text{L}$ |
| $N_{env}$ | number of sub-population testing environments<sup>4</sup> | 100 |
| $N_{pop}$ | population size | 5000 |
| $SD_p$ | standard deviations for basal production rate | 10<sup>-10</sup> |
| $SD_r$ | standard deviations for auto-regulation ratio | 1 |
| $SD_{S_{Th}}$ | standard deviations for signal response threshold | 0.1 |
| $S_{Th_{\text{init}}}$ | initial value for signal response threshold | 5 $\mu\text{L}$ |
| $S_{Th_{\min}}/S_{Th_{\max}}$ | the minimum/maximum signal response threshold | 0.001/20 $\mu\text{M}$ |
| $\lambda_p$ | mutate rate for basal production rate | 0.01 |
| $\lambda_r$ | mutate rate for auto-regulation ratio | 0.01 |
| $\lambda_G$ | parameter used in the zero-truncated Poisson distribution | 0 to 10 |
| $\lambda_{S_{Th}}$ | mutate rate for signal response threshold | 0.01 |
| $p_{\min}/p_{\max}$ | the minimum/maximum basal production rate | $0/2 \times 10^{-8} \mu\text{Ms}^{-1}$ per cell |
| $p_{\text{init}}$ | initial value for basal production rate | $0.5 \times 10^{-8} \mu\text{Ms}^{-1}$ per cell |
| $r_{\min}/r_{\max}$ | the minimum/maximum auto-regulation ratio | 0.01/50 |
| $r_{\text{init}}$ | initial value for auto-regulation ratio | 10 |
| $u$ | signal decay rate | 10<sup>-4</sup> s<sup>-1</sup> |

<sup>1</sup> FU: fitness unit<sup>2</sup> CTE: cooperative testing environment<sup>3</sup> CB: cooperative behavior<sup>4</sup> The local cellular densities are evenly spaced within the range 10<sup>1.5</sup> to 10<sup>5</sup> (cells per $\mu\text{L}$ ).

## Footnotes

1 Note that half of total testing environments are regarded as ‘wrong’ environments since we set N_Th_as the median cellular density across all testing environments.

2 Here, we only consider non-cheats. In other words, the existing cheats in the population pool will not be chosen.

3 The individual genotype’s cooperative investment is simply defined as the number of sub-population testing environments where cooperation is turned on.

